# Comparative Analysis of Orthologous ANK-NLR-WRKY Proteins in Wheat Species: Evolutionary Divergence and Structural Characteristics

**DOI:** 10.64898/2026.11.18.767528

**Authors:** Naledi Mkhize, Eduard Venter

## Abstract

The plant nucleotide-binding leucine-rich repeat (NLR) immune receptors are known to play a crucial role in plant host defence against a variety of biotic stresses. Several NLR proteins contain integrated domains (NLR-ID) like ankyrin (ANK) and WRKY transcription factor domains and are important evolutionary adaptations in plant immunity. They are responsible for the perception of effector molecules from invading pathogens and pests and subsequent activation of plant immune responses. However, the functional roles these ANK-NLR-WRKY proteins play in wheat (*Triticum aestivum*) against various pathogens and pests remain poorly understood. Computational tools employed for comparative sequence and phylogenetic analyses revealed that *TaAdnr1* is highly conserved across diploid (*T*. *urartu*), tetraploid (*T*. *dicoccoides*), and hexaploid (*T*. *aestivum*) wheat species, including its well characterised orthologue YrU1 from *T. urartu*. Three-dimensional (3D) protein structural modelling using AlphaFold2 supported strong conservation in the multiple domains of these NLR orthologues, more specifically within the integrated WRKY domain. This conservation may suggest a strong evolutionary maintenance of TaADNR1. Furthermore, this may also suggest TaADNR1 reflects specificity in effector recognition and regulation. Therefore, the integration of functional genomics studies, sequence analysis, evolutionary analysis and structural modelling possesses the potential to provide new insights into the pleiotropic roles of integrated domain NLRs in wheat.

## Introduction

The WRKY transcription factor (TF) domain consists of the conserved amino acid sequence WRKYGQK at its N-terminus, which is typically around 60 amino acids in length and in some instances includes domain repetitions (Hussain et al., 2024). These domains are known to bind to specific W-box elements ((T/C)TGAC(T/C)) in the promoter regions of target genes, thereby modulating their transcription (Zhang et al., 2020; Zhang & Wang, 2005). The WRKY proteins are classified into three major groups based on the number of WRKY domain sequences and the features of their zinc-finger motifs (Zhao et al., 2023). Group I consists of two WRKY domain sequences, with group II and group III containing only one WRKY domain sequence. Group I and II WRKY proteins are characterised by the presence of C2H2 zinc finger motifs while Group III WRKY proteins contains a C2HC zinc finger motif (Zhao et al., 2023). WRKY proteins are further subdivided into various subgroups based on phylogenetic analyses, which reveal distinct evolutionary pathways and functional specialisation among them (Sinha et al., 2024). The presence of these structures and sequences enables the WRKYs to bind to W-box cis-elements in the target gene promoter (Zhang & Wang, 2005). These structural distinctions within the WRKY family are critical for their varied DNA-binding specificities and diverse regulatory functions across plant species (Sinha et al., 2024).

WRKY transcription factors interact with signalling pathways and coordinate a complex regulatory network essential for plant defence and environmental adaptation (Villano et al., 2020; Wen et al., 2023). These pathways are mediated by various phytohormones and secondary messengers, including salicylic acid and jasmonic acid, which control the plant’s defence mechanisms against pathogens and pests (Zhao et al., 2023; Chen et al., 2019). They modulate the expression of diverse stress-related genes by binding to their cognate W-box elements and initiate or repress gene expression to facilitate adaptive responses. Furthermore, they also interact with other transcription factor families, such as those from the bHLH, NAC, and MYB families, forming complex regulatory networks that regulate plant responses (Finatto et al., 2018). This highlights the pleiotropic roles of WRKY proteins in orchestrating the intricate molecular responses to diverse environmental settings and plant developmental stages (Hussain et al., 2024; Zhang et al., 2020).

Plant nucleotide-binding leucine-rich repeat (NLR) proteins are a vital component of the plant innate immune system for the recognition of pathogen or pest invasion and the subsequent activation of a rapid and robust immune response (Césari, 2018). The architecture of plant NLR proteins consists of a complex mechanism of defence and a comprehensive evolutionary divergence in both monocotyledonous and dicotyledonous plants (Shao et al., 2016). These NLR proteins possess additional domains that provides an evolutionary strategy to detect pathogen invasion. Such NLRs consist of additional non-canonical domains that resemble host proteins that are targets of pathogen effectors and are termed integrated decoy NLRs (ID-NLRs) (Ellis, 2016; Gu et al., 2025). This strategy is a product of processes such as domain fusions, diversification and gene duplication that enable plants to adapt to the evolving pathogens (Césari et al., 2014). Studies have reported that these additional domains are implicated in the recognition of pathogen effectors either directly or indirectly. Following recognition by these intricate IDs, a signalling cascade is activated which results in programmed cell death (PCD) and systemic acquired resistance (SAR) (Marchal et al., 2022; Sarris et al., 2015). The WRKY and zinc-finger BED domains are examples of the IDs that are frequently identified and studied in plants (Kroj et al., 2016) with the diversification of these IDs a prominent evolutionary strategy for NLR diversification (Ellis, 2016).

The WRKY domain-integrated NLR genes play a crucial role in wheat defence by enhancing transcriptional regulation during pathogen and pest attacks. Plant NLRs with WRKYs suggest that pathogen recognition and transcriptional reprogramming are interconnected as many WKRY transcription factors have been implicated in these roles (Anbu et al., 2023; Jin et al., 2022). This enables plants to produce a rapid and precisely regulated defence response within the host (Chen et al., 2019). In wheat, the WRKY integrated NLR orthologs, *TaAdnr1* and *YrU1* were identified from *T. aestivum* and *T. urartu*, respectively (Wang et al., 2020a; Nicolis & Venter, 2018). Studies on *TaAdnr1* and *YrU1* represent defence mechanisms where the integrated WRKY domain likely mediates the transcriptional activation of defence pathways upon pathogen or pest recognition. Knockdown of *TaAdnr1* containing a single WRKY integrated domain resulted in increased susceptibility to the aphid *Diuraphis noxia* (Nicolis & Venter, 2018), with the *YrU1* WRKY linked to stripe rust resistance against *Puccinia striiformis* f. sp. *tritici* (*Pst*) (Wang et al., 2020). The unique structure of *YrU1* was shown to aid in the recognition of pathogenic effectors and thereby activating immune signalling (Wang et al., 2020a). This functional integration of a transcription factor domain directly within an NLR sensor provides a mechanism for the immediate transcriptional modulation upon effector recognition and subsequent initiation of defence responses (Anbu et al., 2023).

The role of integrated WRKY domains into NLR proteins is best studied for *Arabidopsis* RRS1. The integrated WRKY domain of RRS1 allows for the direct recognition of pathogen effectors such as PopP2, which acetylates the integrated WRKY domain, leading to NLR activation and subsequent immune responses (Sarris et al., 2015). This mechanism exemplifies an integrated decoy strategy, where the WRKY domain within RRS1-R mimics a host target of PopP2, thereby converting effector-mediated suppression of host defences into immune activation (Roux et al., 2015). The deletion of the WRKY domain in RRS1 resulted in an auto-active allele, suggesting that this domain actively contributes to maintaining the NLR complex in an inactive state prior to ligand detection (Ma et al., 2018). Upon effector binding, this inhibitory effect is released and enables activation of the associated NLR and subsequent initiation of defence responses (Li et al., 2019). This intricate regulatory mechanism defines sophistication of plant innate immunity where NLR-WRKY fusions offer an alternative strategy for detecting pathogen effectors that specifically target host transcription factors (Chen et al., 2019).

This evolutionary strategy suggests that integrated domains in plant NLRs, including WRKY and others, are crucial for detecting various effectors and cofactors in plant immunity for activation of an immune signal and ultimate defence response. This intricate mechanism enhances the specificity as well as the efficacy of the plant immune response (Dangl et al., 2013). Furthermore, this integration strategy allows for an efficient method for the NLR to directly perceive effector modifications of these host targets and convert the pathogen’s virulence strategy into an immune activation trigger (Adachi et al., 2019). To further our understanding of these roles we investigated the predicted 3D architectures of the different wheat ANK-NLR-WRKY and their evolutionary relationships, to evaluate the possible similarities and differences between them, as well as the potential conserved resistance mechanisms, using molecular biology computational tools.

## Material and Methods

### Wheat ANK-NLR-WRKY protein (*T. aestivum*, *T. dicoccoides* and *T. urartu*) sequence retrieval, domain identification and alignment

The full-length canonical protein sequences of TaADNR1 (*T. aestivum*, AABBDD genome hexaploid wheat), TRIDC5AG050380 (*T. dicoccoides*, AABB genome tetraploid wheat) and YrU1 (*T. urartu*, AA genome diploid wheat and A-genome ancestor for tetra- and hexaploid wheat), were downloaded from EnsemblPlants. The domain boundaries of the three proteins for the NB-ARC and WRKY domains were predicted using InterProScan (Blum et al., 2026) searches against the plant family PF00931 for the NB-ARC domain and PF03106 for the WRKY domain. The coordinates of the domain region of the WRKY domains were extracted and validated by the presence of the conserved WRKYGQK motif. The three full-length wheat ANK–NLR–WRKY proteins were aligned using MAFFT (v7) utilising the L-INS-i algorithm. The default gap opening and extension penalties from MAFFT were used and the alignment was imported to the web-based ESPript 3.0 tool (https://espript.ibcp.fr/ESPript/ESPript/) for visualisation. The parameters selected in the ESPript 3.0 tool included %MultAlin for generation of an alignment with similar residues according to their chemical properties and a default Global score of 0.7 since the proteins are long with multiple domains. The generated ESPript alignment output showed conserved residues based on the similarity thresholds. Finally, the produced ESPript alignment output was used to annotate the different domains (ANK, NB-ARC and LRR domains) as well as the conserved motifs (WRKYGQK and C2HC-type zinc-finger motif) inferred using the InterProScan annotations from the BLAST search outputs.

The isolated WRKY domain sequences from the three wheat proteins and *Arabidopsis* AtWRKY41 and AtWRKY46, each consisting of 60 amino acids, were extracted according to the Pfam WRKY DNA-binding domain (PF03106) based on InterProScan annotations. These WRKY-only sequences for YrU1, TRIDC5AG050380, and TaADNR1 were aligned using MAFFT (v 7) with default parameters and the alignment was visualised using ESPript 3.0. They were chosen for further computational analyses as the phylogenetic analysis indicated that they are the potential WRKY domain progenitors found in YrU1 and TaADNR1.

### Evolutionary relationships of ANK–NLR–WRKY proteins across grasses

### Wheat NLR-WRKY protein sequence retrieval, alignment and construction of phylogenetic tree

The full-length canonical protein sequences consisting of the NB-ARC and WRKY domain from wheat and related grass species were obtained from EnsemblePlants and aligned using MAFFT with the L-INS-i algorithm, which is optimised for accurate alignment of protein sequences containing conserved motifs and variable regions (Katoh & Standley, 2013). ClipKIT v1.4.1 in smart-gap mode was used to trim the MAFFT alignment to reduce alignment noise and poorly aligned regions and to retain phylogenetically informative sites while removing columns dominated by gaps (Steenwyk et al., 2020). The maximum-likelihood phylogenetic tree was constructed using IQ-TREE v2, branch support was assessed by using 1 000 ultrafast bootstrap replicates, and the tree was rooted using the *Arabidopsis* RRS1 sequences as outgroups.

### Phylogenetic analysis and identification of TaADNR1 WRKY transcription factor lineage associated with WRKY domain integration

The complete WRKY transcription factor proteome of *T. urartu* was combined with the WRKY domain sequences of YrU1, TRIDC5AG050380, and TaADNR1. *Arabidopsis* WRKY1, WRKY40, and WRKY55 were included as reference outgroups to anchor the tree relative to well-characterised WRKY lineages. All the *T. urartu* WRKY transcription factors that clustered with YrU1, TRIDC5AG050380, and TaADNR1, were considered as possible candidate donor lineages for WRKY domain integration. The candidate *T. urartu* WRKY transcription factors identified from the clustering were annotated by cross-referencing EnsemblPlants gene IDs with WRKY family classifications, *Arabidopsis* orthologues in the Plant Transcription Factor Database (PlantTFDB) as well as published functional studies.

### AlphaFold wheat (*T. aestivum*, *T. dicoccoides* and *T. urartu*) ANK-NLR-WRKY structure prediction and comparison

The protein sequences of the different wheat ANK-NLR-WRKY proteins (TaADNR1, YrU1, and TRIDC5AG050380) and *Arabidopsis* WRKY proteins AtWRKY41 and AtWRKY46 were submitted to AlphaFold2, v2.3.2, (Jumper et al., 2021) for structural prediction via the Latch.bio platform using the default parameters. From the results of the prediction, the highest-ranked model (ranked_0.pdb) for each protein was used for all the downstream analyses. The per-residue confidence scores (pLDDT) were extracted from the B-factor column of the predicted PDB files using a custom Python script (Supplementary data), utilising the NumPy library. The average pLDDT was calculated for the full-length of each of the protein, as well as the WRKY specific domain according to Supplementary Table 1 (Table S1). The residues were categorised by confidence percentage with very high (>90%), high (70-90%), low (50-70%), and very low (<50%).

For each of the proteins, the model structure with the highest rank was visualised, superimposed, and further analysed using the UCSF ChimeraX (v1.8) software. The pairwise alignment of the structures for the three wheat proteins, and TaADNR1 with AtWRKY41 and AtWRKY46, were performed using the matchmaker command in ChimeraX. The model structures of YrU1 and TRIDC5AG050380 were aligned to TaADNR1 as the reference. ChimeraX was also used to calculate the root-mean-square deviation (RMSD, in Ångströms) and was reported for both the full-length superpositions, as well as the isolated WRKY domains. The WRKY domains were isolated for each of the proteins following superpositions by selecting the defined residue ranges and hiding the non-selected atoms for a focused visual analysis.

## Results

### Wheat ANK-NLR-WRKY protein (*T. aestivum*, *T. dicoccoides* and *T. urartu*) sequence retrieval, domain identification and alignment

The domain architecture of the three ANK–NLR–WRKY proteins from wheat, TaADNR1 (*T. aestivum*), TRIDC5AG050380 (*T. dicoccoides*) and YrU1 (*T. urartu*) revealed that they were highly similar and conserved in the N-terminal ankyrin repeat (ANK) domain and the central NB-ARC domain, with minor amino acid substitutions within the C-terminal leucine-rich repeat (LRR) region and the terminal WRKY DNA-binding domain. The domain boundaries were defined using InterProScan annotations (Table 1).

**Table 1:** The domain architecture of the different wheat ANK-NLR-WRKY proteins.

| **Protein** | **Source** | **Full length (aa)** | **NB-ARC domain (aa)** | **WRKY domain (aa)** |
|---|---|---|---|---|
| **TaADNR1** | *T. aestivum* (Hexaploid) | 1482 | 167 (504-671) | 59 (1333-1392) |
| **TRIDC5AG050380** | *T. dicoccoides* (Tetraploid) | 1482 | 167 (504-671) | 59 (1333-1392) |
| **YrU1** | *T. urartu* (Diploid) | 1576 | 167 (522-689) | 59 (1429-1488) |

The full-length sequence alignments shared 100% amino acid identity in all the ANK and NB-ARC domains for the three proteins (Figure S1) strongly supporting the evolutionary conservation of these regions. The LRR region shared 100% complete identity between TaADNR1 and TRIDC5AG050380. YrU1 exhibited 98.37% identity against both TaADNR1 and TRIDC5AG050380 proteins. Contrastingly, six amino acid substitutions were identified in the LRR region of YrU1 compared to both TaADNR1 and TRIDC5AG050380 that showed 100% identity in this region. The six amino acid substitutions from the similar TaADNR1 and TRIDC5AG050380 compared to the YrU1 sequence were, Ser1154Thr, Gly1199Arg, Ile1205Met, Thr1216Ala, Tyr1223Ala, and Ala1235Asp. The conserved WRKY domain WRKYGQK motif as well as the C2HC-type zinc-finger motif was retained in all the three proteins, which indicated that the DNA-binding functionality should be maintained. There were some amino acid differences between TaADNR1 and TRIDC5AG050380 and YrU1 observed before and after the WRKY domains.

High conservation was observed from the multiple sequence alignments of the isolated WRKY domains of the three wheat proteins, especially on the WRKY core. TaADNR1 and TRIDC5AG050380 shared 100% identity and YrU1 indicated 83.33% identity compared to both proteins (Figure 1a). The WRKYGQK motif was completely conserved in all three proteins which indicated preservation of DNA-binding specificity. In contrast, there were nine amino acid substitutions observed in the regions downstream of the WRKY motif in YrU1 compared to TaADNR1 and TRIDC5AG050380. These substitutions included changes Thr14Ile, Met15Ile, Leu35Pro, Ala36Val, Asn45Thr, His54Tyr, and Asn55His. There was only one amino acid substitution observed upstream of the WRKY motif at position 1 (Asp1Glu). Importantly, there were no substitutions observed in WRKYGQK motif as well as the conserved zinc-finger motif in all the three proteins (Figure 1a, Figure S1).

**Figure 1:**
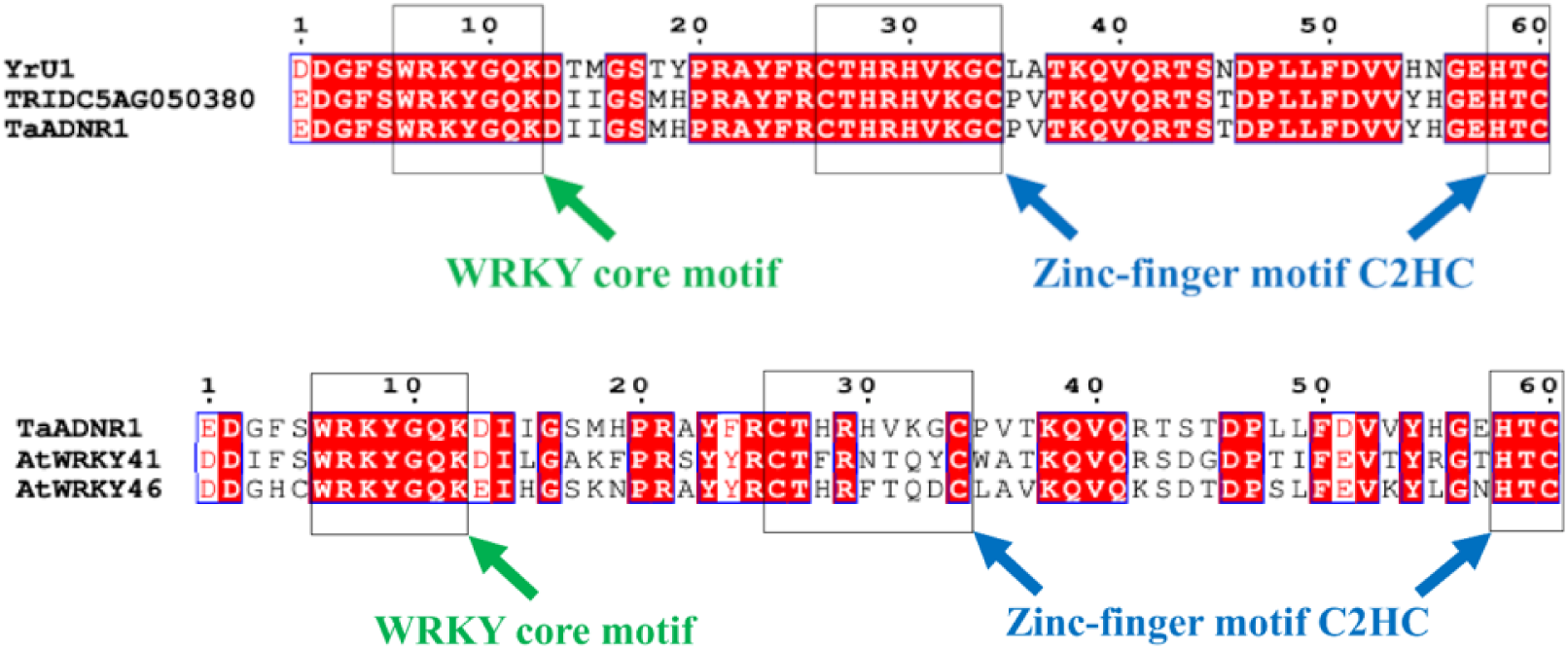
**A**, The WRKY domain-only multiple sequence alignments isolated from the ANK–NLR–WRKY protein sequences of TaADNR1, TRIDC5AG050380 and YrU1 (*T*. *aestivum*, *T*. *dicoccoides* and *T*. *urartu*)The WRKY domain-only multiple sequence alignments of the isolated WRKY regions from TaADNR1, AtWRKY41 and AtWRK46.Amino acid substitutions are present upstream and downstream the WRKY core, including the zinc-finger motifs in AtWRKY41 and AtWRKY46. There were no amino acid substitutions observed within the WRKY core for all the proteins.

The sequence alignment between TaADNR1 and AtWRKY41 shared 60% amino acid identity, TaADNR1 and AtWRKY46 shared 61.67% amino acid identity, and AtWRKY41 and AtWRKY46 shared a 66.67% identity (Figure 1b). The alignments of the isolated WRKY domain of TaADNR1 and both the *Arabidopsis* WRKY protein domains revealed that the WRKY core was conserved. There were twenty-four amino acid substitutions for TaADNR1 against AtWRKY41, twenty-three between TaADNR1 and AtWRKY46, and twenty amino acid substitutions between AtWRKY41and AtWRKY46. These substitutions were observed in the region upstream and downstream the WRKY core. Notably, substitutions were also observed in the region within the Zinc-finger motif between the three proteins.

There were sixteen amino acid substitutions between TaADNR1 and AtWRKY41 that showed the most important polarity and charge changes, as well as broad physicochemical class changes (Table S1). There were eight amino acid substitutions that retained the same broad class changes. The sixteen amino acid substitutions that showed the most important polarity and charge changes included Ser17Ala and Thr45Gly, changing from polar uncharged to non-polar, His19Phe and His28Phe, changing from polar-ionisable to non-polar. There were six amino acid substitutions that exhibited changes from non-polar to polar uncharged and these included Ala22Ser, Phe24Tyr, Val31Thr, Gly33Tyr, Leu48Thr and Val53Thr. Single amino acid substitutions were observed for changes from non-polar to polar positively charged (Met18Lys), polar uncharged to polar negatively charged (Ser44Asp), polar positively charged to polar uncharged (Lys32Gln), polar ionisable to polar positively charged (His55Arg) and polar negatively charged to polar uncharged (Glu57Thr).

The alignment between TaADNR1 and AtWRKY46 indicated fifteen amino acid substitutions with the most important polarity and charge changes, and broad physicochemical class changes (Table S2). There were eight amino acid substitutions that retained the same broad class changes. The fifteen amino acid substitutions that indicated the most significant polarity and charge changes included Thr37Val, changing from polar uncharged to non-polar, His28Phe and His55Leu, changing from polar-ionisable to non-polar. There were three amino acid substitutions that displayed substitutions from non-polar to polar uncharged and these included Phe24Tyr, Val31Thr and Leu48Ser. Substitutions from non-polar to polar ionisable and non-polar to polar positively charged included Phe4His and Ile15His, and Met18Lys and Val53Lys, respectively. Single amino acid substitutions were observed for changes from non-polar to polar negatively charged (Gly33Asp), polar uncharged to polar negatively charged (Ser44Asp), polar positively charged to polar uncharged (Lys32Gln), polar ionisable to polar uncharged (His19Asn) and polar negatively charged to polar uncharged (Glu57Asn).

The AtWRKY41 and AtWRKY46 alignment showed fifteen amino acid substitutions that were non-identical with significant polarity and charge changes (Table S3). These included Ser22Ala, changing from polar uncharged to non-polar and Ala17Ser, Phe19Asn and Gly45Thr, changing from non-polar to polar uncharged. There were three amino acid substitutions that displayed substitutions from non-polar to polar ionisable and these included Phe4His, Leu15His and Phe28His. Single amino acid substitutions were observed for changes from polar uncharged to polar positively charged (Thr53Lys) and polar positively charged to non-polar (Arg55Leu).

### Phylogenetic relationships between NLR-WRKY proteins across grasses and placement of the integrated WRKY domain

The evolutionary relationships of NLR-WRKY proteins across monocot grass species included *T*. *aestivum* (TaADNR1), *T*. *dicoccoides* (TRIDC5AG050380), *T. urartu* (YrU1), and homologues from *Aegilops*, *Oryza*, and *Brachypodium* with the resulting topology indicating TaADNR1, TRIDC5AG050380 and YrU1 forming a strongly supported (100 %) monophyletic clade (Figure 2). The WRKY-only phylogenetic tree was constructed using the complete *T. urartu* WRKY transcription factor proteome isolated from EnsemblPlants, the WRKY domains derived from YrU1, TRIDC5AG050380, and TaADNR1 full-length sequences, and the *Arabidopsis* WRKY1, WRKY40, and WRKY55 as outgroups (Figure 3). This was to identify the likely WRKY transcription factor lineage that contributed the integrated WRKY domain into the wheat NRL protein. The WRKY domains from YrU1, TRIDC5AG050380, and TaADNR1 clustered together in a well-supported clade (≥ 99 %) with YrU1 basal to TRIDC5AG050380, and TaADNR1 (Figure 3). This confirmed the shared evolutionary origin of these proteins with YrU1 as ancestral to TRIDC5AG050380 and TaADNR1. This clade grouped well with five *T. urartu* WRKY transcription factors that included WRKY41 and WRKY46 suggesting that these WRKY’s represent the closest relatives of the WRKY donor lineage. A BLAST search against *Arabidopsis* WRKY proteins from the Plant Transcription Factor Database (PlantTFDB) and The Arabidopsis Information Resource (TAIR) confirmed the six *T. urartu* WRKY transcription factors were similar to the WRKY41- and WRKY46-like proteins (Figure S2, Table S4). Four of the candidates (TuG1812G0100004483.01, TuG1812G0300002773.01, TuG1812S0003165700.01, TuG1812G0200001696.01) showed similarity to WRKY41-like protein, with TuG1812G0500002157.01 exhibiting similarity to the WRKY46-like protein. These WRKY41-and WRKY46-like sequences showed strong sequence similarity (E-values ≤ 10⁻³¹, Table S4), high bootstrap support (76–100 %, Figure 3), and clustered tightly with the integrated WRKY domains to support a close evolutionary relationship.

**Figure 2:**
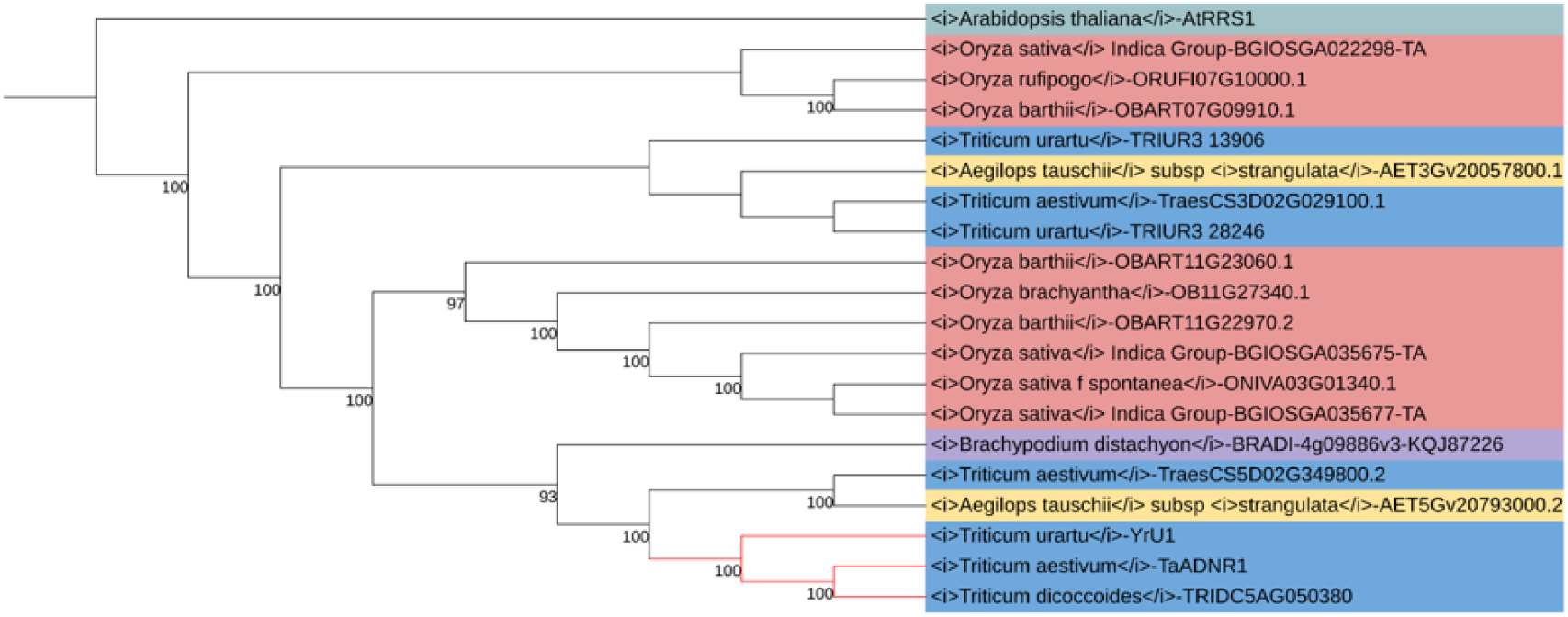
The phylogenetic relationships of the wheat NLR-WRKY proteins

**Figure 3:**
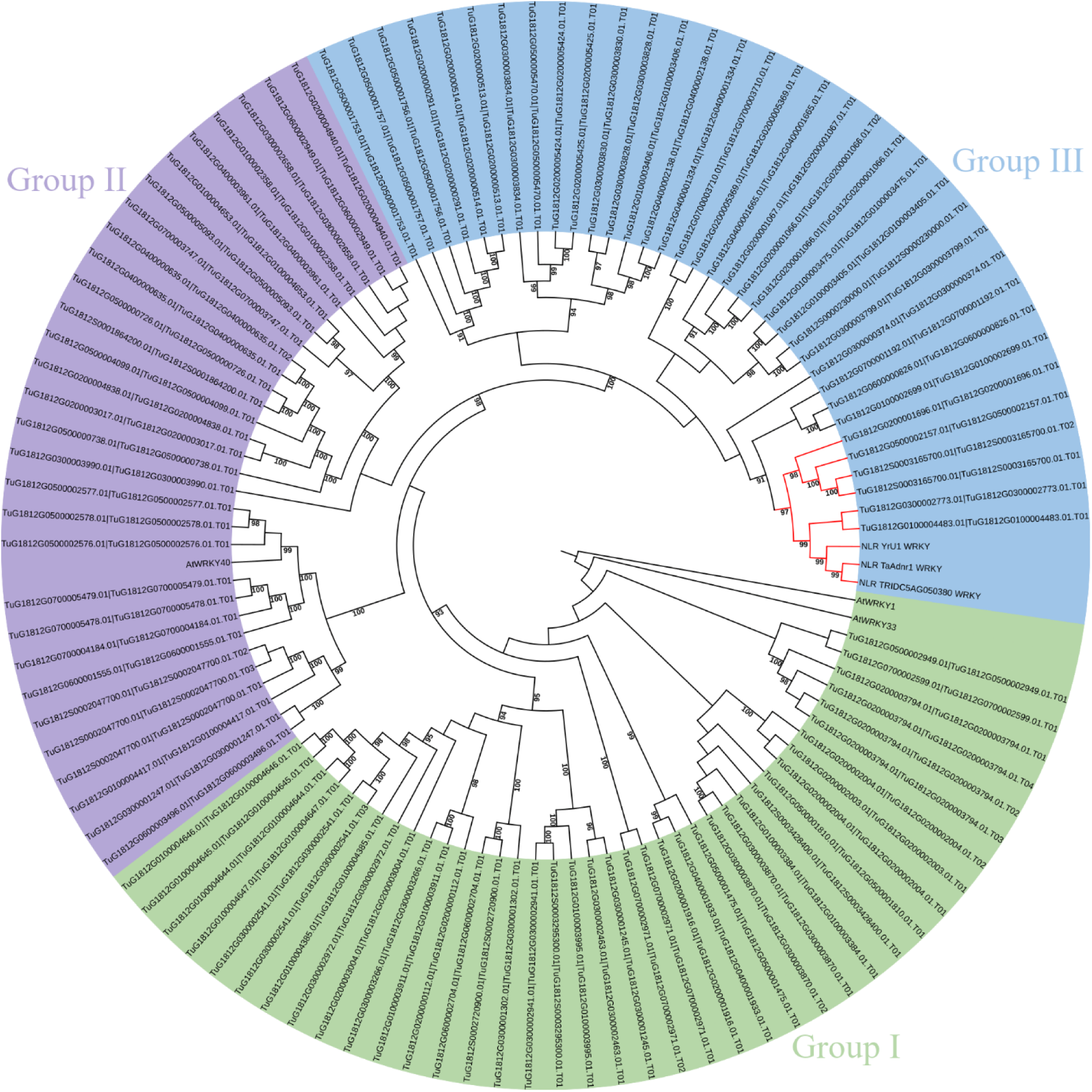
The WRKY-only phylogenetic analysis for the identification of the likely donor lineage of the integrated WRKY domain

### AlphaFold2 structure predictions, comparison and analysis of wheat ANK-NLR-WRKY full-length proteins and WRKY domain

The AlphaFold2 structure predictions generated quality structural models for the three full-length ANK-NLR-WRKY proteins (Figure 4, Table S5). The full-length model for TaADNR1 and TRIDC5AG050380 showed a high average pLDDT scores of 71.91 and 71.64 (Figure 4A and 4C), respectively. YrU1 indicated a moderate average pLDDT score of 68.32 (Figure 4B). The average pLDDT scores for the WRKY specific domain of all three proteins was high with 82.27, 81.61 and 78.48 for TaADNR1, TRIDC5AG050380 and YrU1, respectively. Notably, both TaADNR1 and TRIDC5AG050380 WRKY domains exhibited pLDDT scores above 81.

**Figure 4:**
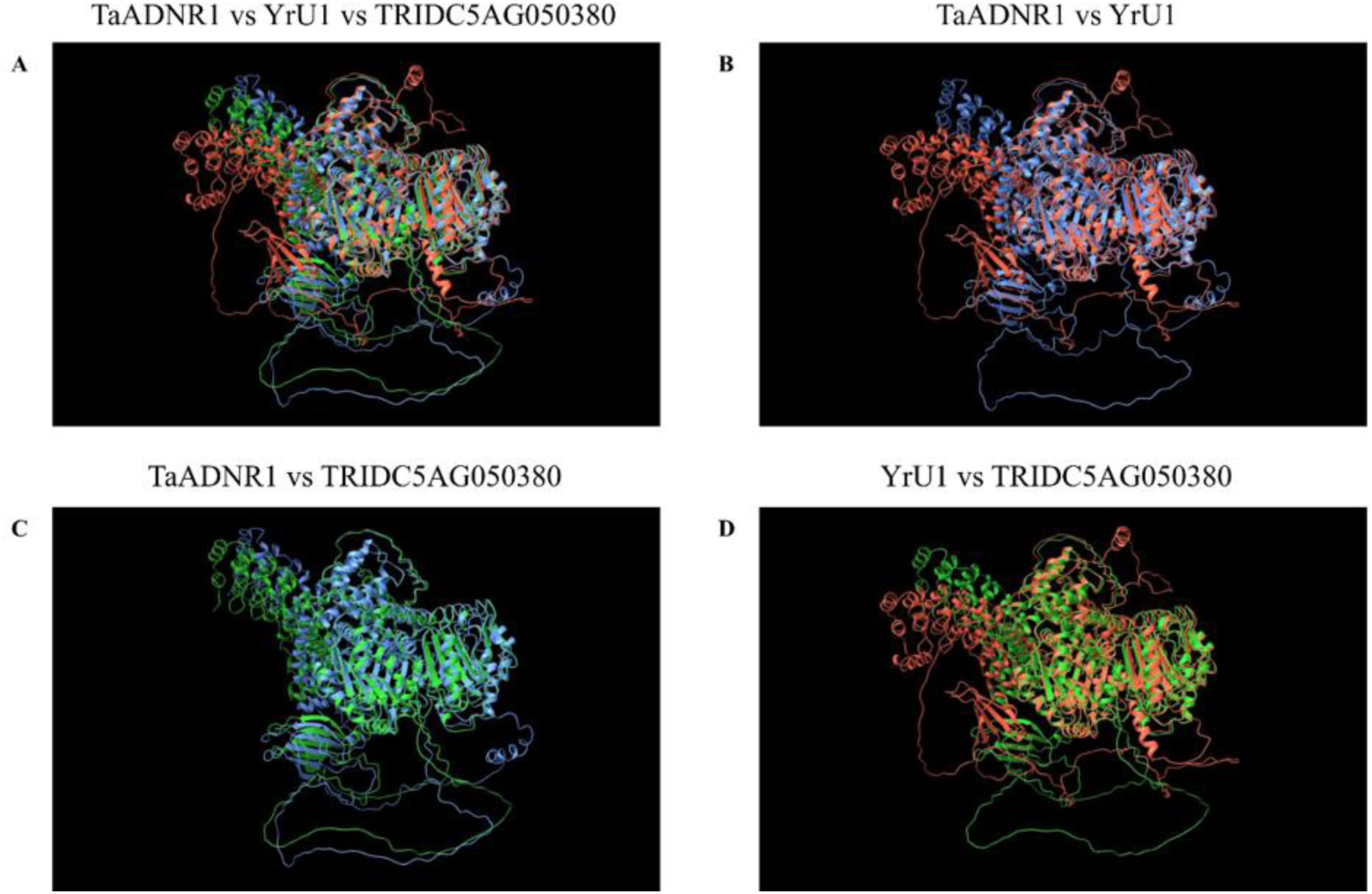
Alphafold2 predicted structural alignment of full-length ANK-NLR-WRKY proteins

**Figure 5:**
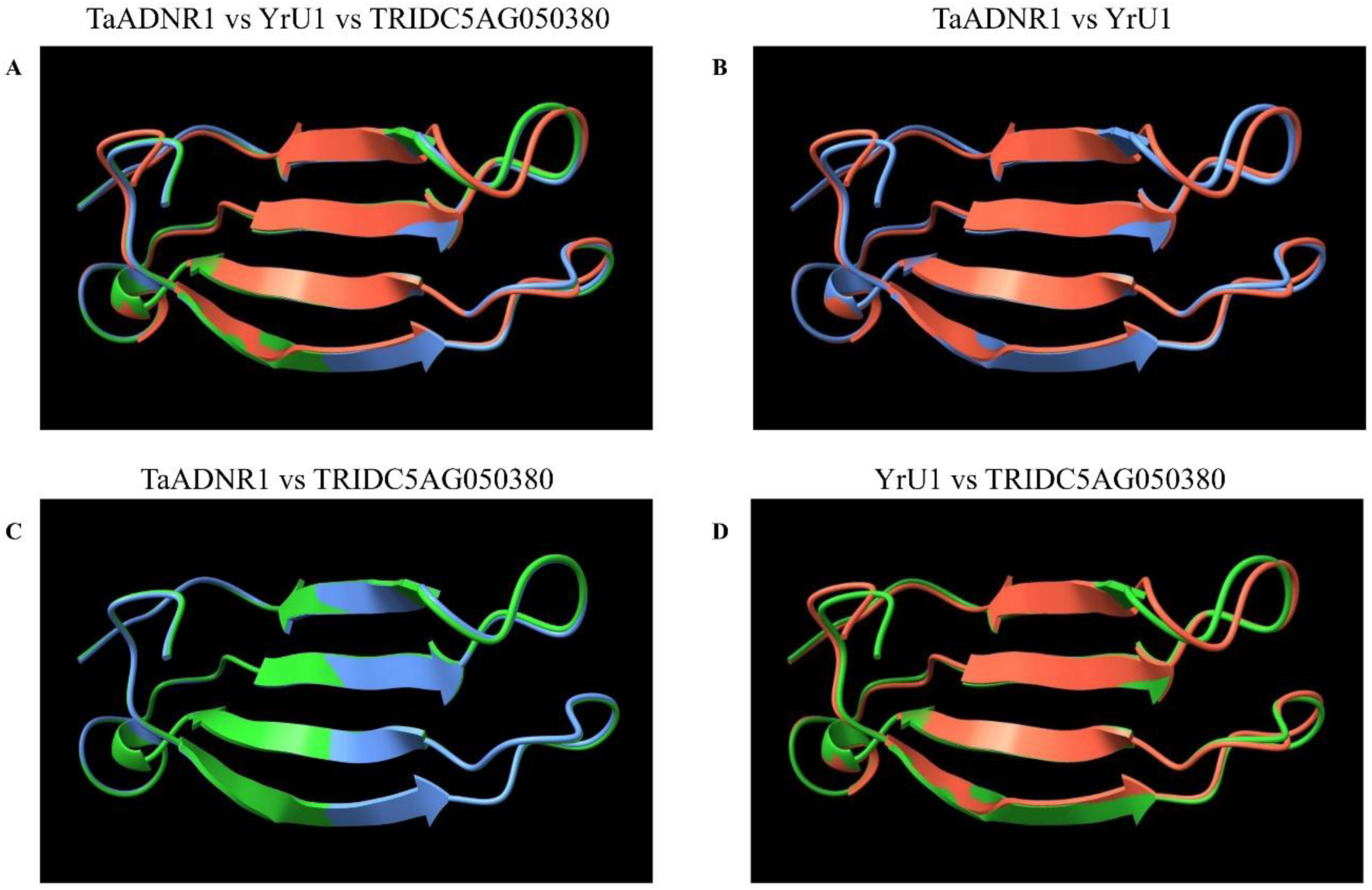
The structural alignment of the isolated WRKY domains of the wheat (*T*. *aestivum*, *T*. *urartu* and *T*. *dicoccoides*) ANK-NLR-WRKY proteins

The structural superposition revealed a high degree of conservation across the three proteins (Figure 4, Table S5). The three ANK-NLR-WRKY full-length proteins showed high similarity with pairwise RMSD values below 0.75 Å (Figure 4A, Table S5). The structural superposition between TaADNR1 and YrU1 indicated a low RMSD of 0.746 (Figure 4B, Table S5), while TaADNR1 and TRIDC5AG050380 exhibited the lowest RMSD of 0.424 (Figure 4B, Table S5). The conservation was even more pronounced within the WRKY domain. Notably, the WRKY domains of TaADNR1 and TRIDC5AG050380 were nearly indistinguishable, with an RMSD of only 0.130 Å (Figure 4C, Table S5). The WRKY domain of the more distantly related YrU1 also maintained very high similarity to TaADNR1 RMSD 0.560 Å (Figure 4B, Table S5). A similar trend was observed in the high-confidence residue (70-90 %) score, with the full-length proteins indicating reduced low-confidence scores of 53.6 %, 53.9 % and 52.1 % for TaADNR1, TRIDC5AG050380 and YrU1, respectively (Figure 6A-C, Figure S2, Table S5).

**Figure 6:**
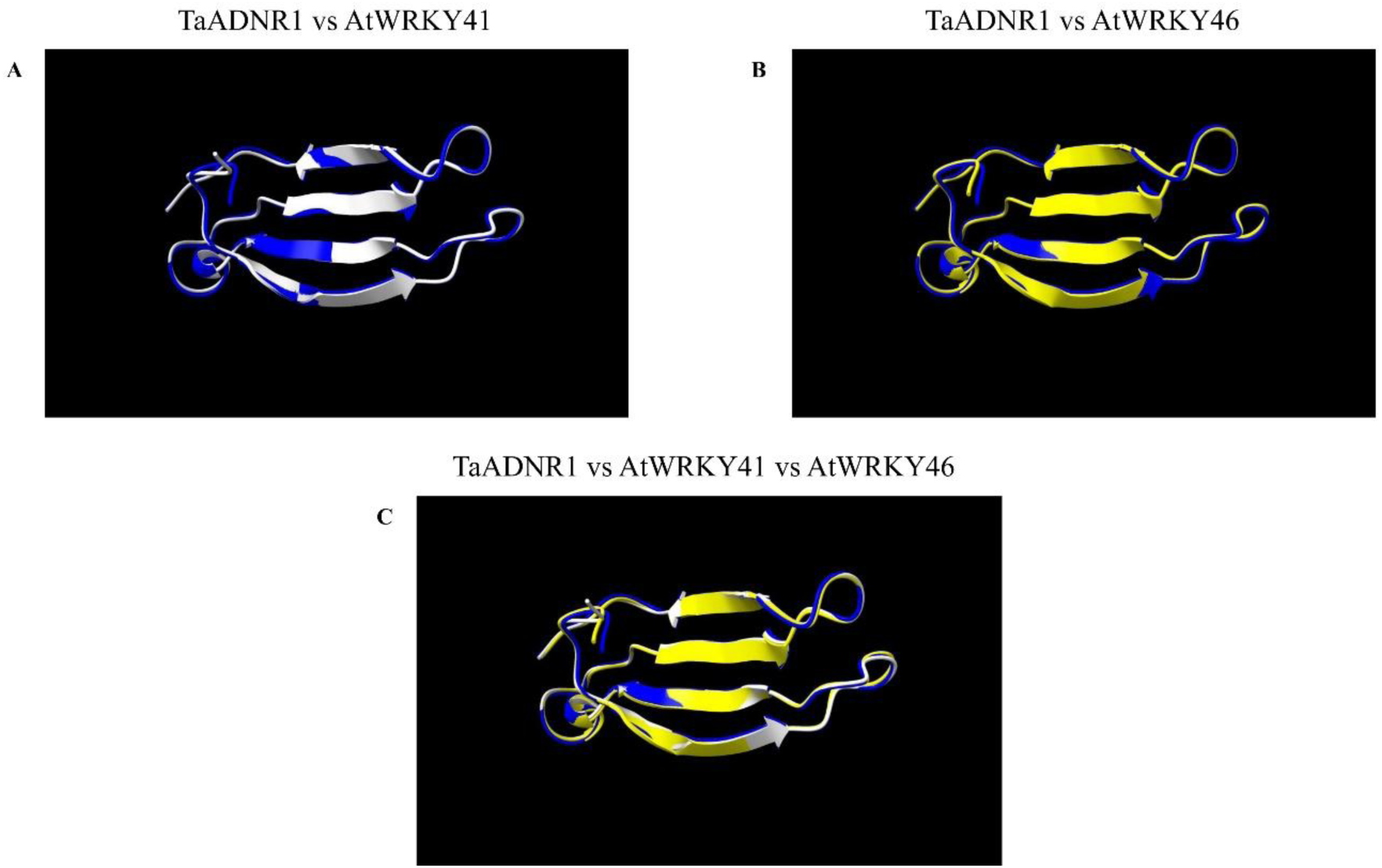
The structural alignment of the isolated WRKY domains of the wheat (*T. aestivum*) TaADNR1 WRKY domains against the *A. thaliana* AtWRKY41 and AtWRKY46 proteins. The protein structures are coloured by chain with TaADNR1 WRKY domain coloured blue, AtWRKY41 coloured white and AtWRKY46 coloured yellow. **A** and **B**, Pairwise alignments of TaADNR1 against AtWRKY41 and AtWRKY46, respectively. **C**, Superposition of all three isolated WRKY domains from the three proteins. The near-identical overlap in **A**, with a RMSD of 0.360 Å highlights the high degree of structural conservation with significant differences between TaADNR1 and AtWRKY41 (RMSD < 0.5 Å). This emphasizes that these domain regions adopt strong three-dimensional structural folds regardless of the differences in the amino acid substitutions between the two protein regions.

Compared to the full-length protein, the WRKY specific domain exhibited an increased high-confidence residue score, with higher percentage scores ranging between 76.7-81.7 % (Figure 5, Table S5). Contrastingly, the WRKY-only domain of TaADNR1 and TRIDC5AG050380 both exhibited an increased high-confidence percentage (80 and 81 %, respectively) compared to YrU1 (76.7 %) (Table S5). A reduced low-confidence percentage was similar in both TaADNR1 and TRIDC5AG050380 (6.7 %) compared to the increased low-confidence percentage in YrU1 (18.3 %). Therefore, both TaADNR1 and TRIDC5AG050380 exhibited good low-confidence as well as good high-confidence percentages in the WRKY domain, confirming the high structural prediction for this region. This highlights the high degree of structural similarity and conservation between the two protein domains from the different *Triticum* species.

### AlphaFold2 structure predictions, comparison and analysis of TaADNR1 and AtWRKY41 and AtWRKy46

The AlphaFold predicted structure comparisons of the TaADNR1 isolated WRKY integrated region as well as the *Arabidopsis* AtWRKY41 and AtWRKY46 isolated WRKY region exhibited a high degree of structural conservation between the three proteins regardless of the moderate sequence divergence (Figure 6, Table S6). The average pLDDT for TaADNR1, AtWRKY41 and AtWRKY46 were 82.27, 93.71 and 92.34, respectively (Table S6). This indicated a high confidence in the prediction of the structure with significantly high confidence for both AtWRKY41 and AtWRKY46. The AtWRKY41 protein exhibited a significantly larger high-confidence residue percentage of 88.3% for 53 residues (>90%), while TaADNR1 exhibited a high-confidence residue percentage of 80% for 48 residues (70-90%). The TaADNR1 and AtWRKY41 structural superpositions of the isolated WRKY region indicated a near-identical fold with a RMSD of 0.360 Å, while TaADNR1 and AtWRKY46 exhibited a RMSD of 0.378 Å. This observation demonstrated high similarity in the predicted three-dimensional structure of the compared WRKY regions.

## Discussion

This is the first study that provides a comparative structural analysis of three ANK-NLR-WRKY immune receptor proteins across the wheat diploid, tetraploid and hexaploid species. The analysis of both the full-length and the isolated WRKY domain alignments show how these ANK–NLR–WRKY proteins have evolved in wheat and its relatives. The high degree of similarity and conservation of both the ANK and NB-ARC domains in TaADNR1, TRIDC5AG050380 and YrU1 reflects a strong indication that the structure of these domains are essential for functioning of these immune proteins and do not seem to change significantly over the evolutionary course of time (Mosavi et al., 2004; Steele et al., 2019). This further supports that alteration or disruption of these domains will compromise the functioning of the NLR protein as they hold important roles including NLR activation and downstream defence responses (Takken et al., 2006; Tameling et al., 2006; Wang et al., 2020b).

### Multiple sequence alignments reveal high conservation in the wheat ANK-NLR-WRKYs and WRKY specific domains

The small sequence variations that are present in both the LRR and WRKY domains suggest that these domains are important in enabling the toleration of small changes that may contribute to various adaptations for the functioning of the protein. The LRR domain was recently categorised further, consisting of the highly conserved segment (HCS) and the highly variable segment (HVS) (Liu et al., 2022). YrU1 shows variations within the HVS of its LRR domain compared to both TaADNR1 and TRIDC5AG050380. Such variations are attributed to shaping the functional specificity of the NLR in recognising specific pathogens under positive selection pressures. In contrast, these variations allow the LRR domain to harbour dual functions, including the auto-inhibition of the NLR as well as effector recognition (Anbu et al., 2023).

The wheat WRKY-only alignment shows strict conservation of the WRKYGQK motif as well as the zinc-finger-like motif across all three proteins and confirms the preservation of the core DNA-binding functionality of these proteins. These conserved motifs indicate that they all belong to the group III class of WRKY transcription factors, with the WRKY domains consisting of the C2-HC zinc finger motif and a sequence pattern C-X5-8-C-X25-28-H-X1-2-C (Grzechowiak, 2014). Specifically, the conserved pattern observed for the three proteins is C-X7-C-X23-H-X1-C. These motifs are directly involved in the interaction with the W-box elements in target promoters and are crucial for transcriptional regulation of plant defence-related genes (Bakshi and Oelmüller, 2014). In *Arabidopsis*, the acetylation of the lysine residues within the WRKYGQK domain disrupted the functional activity of RRS1 as well as the RPS4/RRS1 complex for W-box DNA interaction (Sarris et al., 2015). Therefore, the evolutionary conservation of the WRKY core and the zinc-finger motif is crucial for maintaining the high affinity DNA-binding activity.

Nine out of the ten amino acid substitutions identified in YrU1 compared with both TaADNR1 and TRIDC5AG050380 occur mostly downstream of the WRKY core and involve changes in amino acid physicochemical properties. For example, computational analysis suggests that amino acid substitutions from threonine to isoleucine (Thr14Ile) and threonine to methionine (Thr18Met), in YrU1 to TaADNR1 and TRIDC5AG050380, represents non-conservative substitutions that shift from a polar uncharged residue to a non-polar hydrophobic residue. These types of replacements may destabilise a protein’s structure because the introduction of a hydrophobic residue is unfavourable in aqueous environments, while a polar residue could form stable hydrogen bonds with water (Wolynes, 1995). Ultimately, these substitutions may potentially present an increasing flexibility or hydrogen-binding capacity (Abuzaid et al., 2024). Similarly, substitutions Ala36Val and Met15Ile preserves hydrophobicity as they are non-polar. However, the loss of the larger sulphur-containing side chain may impact amino acid local packing in TaADNR1 and TRIDC5AG050380. The Leu35Pro substitution replaces a flexible hydrophobic residue with a more rigid amino acid which may potentially influence the local secondary structure of the domain (Pahuja et al., 2007). Further substitutions such as Tyr19His, His4Tyr and Asn55His, affect the amino acid’s aromatic and charged properties. This has implication in influencing the electrostatic interactions or pH-dependent binding properties. These changes do not necessarily affect the residues that are directly involved in DNA binding and zinc-finger conformations. Indeed, the Alphafold2 modeling supports very limited changes between the WRKY domains of these three proteins. This suggests that the WRKY domain maintains its transcriptional activity and presents regulatory differences or specificity in its interactions (Phukan et al., 2016). Therefore, YrU1 seems to present a diverged ancestral form that is still functional, while TaADNR1 and TRIDC5AG050380 show complete similarity in identity in their WRKY domains. In contrast, YrU1 may display regulatory differences that may contribute to species-specific immune responses, while TaADNR1 and TRIDC5AG050380 may show broad spectrum species recognition in the immune responses (Wang eet al., 2020a). These results indicate that functional activity should be intact. Therefore, the small variations observed in the YrU1 WRKY sequence suggests that they may not negatively affect the overall functioning of the WRKY and may suggest possible differences in gene regulation rather than complete functional divergence.

The amino acid sequence alignment between TaADNR1, AtWRKY41 and AtWRKY46 shows conservation of the WRKYGQK core motif, including the zinc-finger-like motif between wheat and *Arabidopsis*. This observation supports the preservation of the DNA-binding property (Brand et al., 2013; Kim et al., 2024). The substitutions with potential impact on WRKY functioning between TaAdnr1 and AtWRKY41 included His28Phe, Lys32Gln and Gly33Tyr. These substitutions were observed within the zinc-finger core and include changes such as loss of a positive charge (Lys32Gln) and a change from a flexible to a bulkier aromatic amino acid (Gly33Tyr). Furthermore, substitutions outside both the WRKY core and zinc-finger motif included Met18Lys and His55Arg that present introduction of a positive charge, Ser44Asp and Glu57Thr that present local charge creation and removal, and lastly, Ser17Arg and Thr45Gly that represents polar to non-polar changes that can remove hydrogen bonding. Potential impactful substitutions between TaADNR1 and WRKY46 within the WRKY core included Gly33Asp, His28Phe and Lys22Gln. These substitutions may present changes such as loss of ionisation. Substitutions outside both the WRKY core and zinc-finger motif included Val53Lys and Met18Lys that introduces a positive charge, Phe4His and Ile15His that introduces pH sensitivity, Ser44Asp and Glu57Ans that may present electrostatic changes, and lastly, Thr37Val that represents polar to non-polar change. There was only one potentially impactful amino acid substitution between AtWRKY41 and AtWRKY46 within the zinc-finger motif where an ionisable amino acid (Phe28His) was introduced. The other substitutions observed outside the zinc finger motif included Arg55Leu and Thr53Lys that introduced loss of a positive charge and addition of pH sensitivity, respectively. Furthermore, substitutions in Thr53Lys represented an introduction of a positive charge, while Ser22Ala introduced a direct polar-to-non-polar change, and Phe19Asn and Gly45Thr introduced hydrophobic to polar changes that may possibly alter the hydrogen-bonding surfaces. Therefore, these substitutions are suggested to introduce changes in the local hydration, hydrogen bonding, electrostatic potential or packing, without necessarily affecting the folding of the WRKY domain, DNA binding, effector recognition and function of the NLR protein (Kim et al., 2024; Pandey et al., 2018; Mukhi et al., 2021).

### Evolutionary relationships of NLR–WRKY proteins across grass species

The integration of WRKY transcription factor domains into plant NLR’s is a significant evolutionary event in plant immunity as well as other important plant processes in response to environmental stimuli (Mukhtar, 2013). In the present study, we further investigated the evolutionary history of the TaADNR1 from the hexaploid wheat and the origin of the WRKY domain that was integrated into this ANK-NLR-WRKY proteins. The NLR–WRKY phylogenetic tree constructed using NLR’s from different grass species, which consist of the WRKY integrated domain, indicated that TaADNR1, TRIDC5AG050380 and YrU1 (from *T. aestivum*, *T. urartu* and *T. dicoccoides*, respectively) cluster together and form a robust monophyletic clade. This observation supports its origin from a common ancestral gene. The high conservation displayed in the domain architecture in these proteins across the diploid, tetraploid, and hexaploid wheat indicates that the WRKY integration event occurred before polyploidisation of wheat from a distant ancestor and has since been retained. This conservation suggests that the evolutionary integration of the WRKY domain provides an added advantage in the functioning of the NLR, which is consistent with the proposed integrated decoy model for perception of pathogen derived effector molecules (Césari et al., 2014).

Initial studies on TaADNR1 indicated that the most probable WRKY domain donor was the TaWRKY50 transcription factor from group III (Nicolis and Venter, 2018). At the time of this study the *T. urartu* and *T. dicoccoides* genomes were not fully available for comparison when the *T. aestivum* and related grass species’ proteomes were queried. The divergence, 79.7%, over the 69 amino acid WRKY domain raised the question why such a large amount of change took place in such a short timespan (Nicolis and Venter, 2018). The additional information generated through the extended phylogenetic analysis in this study updates this to recognise that the WRKY domain found on TaADNR1, TRIDC5AG050380 and YrU1 originated from a common ancestor and increases our understanding of the evolutionary history of TaADNR1. Similar to Wang et al. (2020), we postulated that the donor of the integrated WRKY domain either contributed to *T. urartu* or possibly *T. dicoccoides*. The phylogenetic tree containing WRKY transcription factors from *T*. *aestivum*, *T*. *dicoccoides* and *T*. *urartu* showed that TaADNR1, TRIDC5AG050380, and YrU1 clustered specifically with WRKYs from *T*. *urartu*. Resolving the tree to focus on *T. urartu* WRKY transcription factors supported a shared an evolutionary origin for the integrated WRKY domains of TaADNR1, TRIDC5AG050380 and YrU1, indicating that TaWRKY41 from *T. urartu* as the closest extant relative of the donor lineage. This narrows the donor contribution to already have occurred in *T. urartu* with subsequent contribution to *T. dicoccoides* and *T. aestivum*.

The six *T. urartu* WRKY candidates (TuG1812G0100004483.01, TuG1812G0300002773.01, TuG1812S0003165700.01, TuG1812G0200001696.01) shared high homology to a WRKY41-like transcription factor and one candidate (TuG1812G0500002157.01) showed homology to WRKY46-like transcription factor similarity to *Arabidopsis* RRS1. Their high sequence similarity with the integrated domains and close phylogenetic clustering further supported the evolutionary relationship with the donor lineage. WRKY41 and WRKY46 are both known regulators of defence signalling, salicylic acid-mediated immunity, and stress responses, making them biologically plausible targets for pathogen effector manipulation (Higashi et al., 2008; Hu et al., 2011; Verk et al., 2011; Ding et al., 2015). Indeed, the AtWRKY41 integrated onto RRS1 recognises AvrRps4 from *Pseudomonas syringae* in *Arabidopsis* (Sarris et al., 2016). This effector trapping leads to the initiation of a defence response against the invading pathogen, which suggests a similar role for integrated WRKY domain on TaADNR1, YRU1 and TRIDC5AG050380.

Collectively, these findings support a model in which a WRKY41-like transcription factor was integrated into an ancestral NLR gene in a diploid wheat progenitor, giving rise to the ANK–NLR–WRKY lineage conserved across wheat evolution. This work advances our understanding of the molecular origins of wheat immune receptors and provides a framework for exploring the functional consequences of WRKY integration in resistance to aphid and fungal pathogens.

### AlphaFold2 structure predictions of wheat NLR-WRKY proteins and WRKY-specific domains

The AlphaFold2 structure predictions generated high-quality structural models for the three wheat ANK-NLR-WRKY proteins. The full-length models exhibited moderate average pLDDT scores (68.32-71.91), a characteristic feature of large, multi-domain intracellular immune receptors which often contain flexible linkers. Shorter proteins generally display higher pLDDT scores for good predictions, and these scores tend to decrease with increasing amino acid length (Ilzhöfer et al., 2022). Prediction confidence, as indicated by pLDDT scores, tends to decline for proteins longer than approximately 1 000 amino acids (Abbas et al., 2023). However, protein sequences with approximately 1 500 aa report moderate to good accuracy in the predictions as well as good confidence in the backbone fold. These scores represent the characteristic nature of large multidomain immune receptors that are often known to consist of flexible linkers between multiple domains in a protein (Jumper et al., 2021; Huang et al., 2023; Hetmann et al., 2024). Therefore, for both TaADNR1 and TRIDC5AG050380 with similar amino acid lengths, the pLDDT scores of 71.91 and 71.64 indicate a moderate to high confidence in the accuracy of the predicted structures. The lower pLDDT score in YrU1 may be attributed to slightly longer amino acid length. These observations are consistent with the fact that there is flexibility in the NLR conformation during activation or inactivation, in the regions where the interacting domains link together (Ohto, 2022).

The isolated WRKY domain regions of the proteins were predicted with significantly higher average pLDDT scores (78.48-82.27). The average pLDDT scores for the WRKY domain specific region of all three proteins was high with 82.27, 81.61 and 78.48 for TaADNR1, TRIDC5AG050380 and YrU1, respectively. Although the scores were not above 90, which may be comparable to experimental resolutions, these high scores, especially for TaADNR1 and TRIDC5AG050380, could indicate a reasonable structural prediction. The high-confidence prediction of the WRKY domains supports that its role in DNA binding is conserved and intact for different NLRs with minimal differences that may regulate distinct signalling mechanisms (Sarris et al., 2016; Anbu et al., 2023).

The integrated WRKY domain of TaADNR1 exhibits a highly conserved 3D conformation compared with the *Arabidopsis* WRKY41 and WRKY46 domains. The average pLDDT scores for the WRKY domain specific region of all three proteins was exceptionally high with 82.27, 93.71 and 92.34 for TaADNR1, AtWRKY41 and AtWRKY46, respectively. Notably, both AtWRKY41 and AtWRKY46 exhibited high-confidence residue percentages with 88.3% (53 residues at > 90%) and 78.7 for (48 residues at > 90%), respectively. These observations indicate that both the AtWRKY41 and AtWRKY46 WRKY domain region may be comparable to experimental resolutions. The low RMSD values of 0.360 Å for AtWRKY41 and 0.360 Å for AtWRKY46 confirm a highly conserved conformation. Despite the moderate degree of amino acid substitutions between the three proteins, the overall 3D conformation was retained. The AtWRKY41 and AtWRKY46 proteins have been previously reported in multiple studies for their contribution in plant defence against various pathogens (Pandey et al., 2018). The study on the structural analysis and conservation of the WRKY domain of the *Solanum lycopersicum* SlWRKY3 and SlWRKY4 revealed high similarity in the structural superposition, with RMSD values less than 0.25 Å and several amino acid substitutions (Aamir et al., 2017). These observations may suggest the possibility that TaADNR1 and both the *Arabidopsis* AtWRKY41 and AtWRKY46 may share the functional ability to recognise pathogen or pest effectors during defence signalling, while exhibiting minor amino acid substitutions. Taken together, the TaADNR1 WRKY integrated domain shares structural conservation across the examined integrated WRKYs from *T. dicoccoides* and *T. urartu*, including closely related *A. thaliana* WRKY proteins that play a role in plant defence.

Pairwise superposition of the WRKY domains revealed a high degree of conservation across the three wheat orthologs. The folds of the full-length proteins were highly similar, with all pairwise RMSD values below 0.75 Å .This suggests that these proteins not only have an identical topology but also share similarities in atomic bond lengths, bond angles, and overall spatial arrangements of their atoms. This high degree of overall structural conservation is notable given the evolutionary divergence between the diploid and polyploid wheat species and suggests strong maintenance of function on the overall NLR-WRKY protein. This high conservation level further supports conservation of function when considering that the NLR family is the largest family within the wheat genome with extensive diversification of its members (Andersen et al., 2020; Walkowiak et al., 2020; Jumper et al., 2021).

A high level of conservation was observed when comparing the isolated WRKY domains in the three proteins. The WRKY domains of TaADNR1 from the hexaploid wheat and its tetraploid progenitor TRIDC5AG050380 were found to be nearly indistinguishable with an RMSD of 0.130 Å. This observation suggests that the 3D fold of this domain has been strictly preserved after the polyploidisation events that gave rise to hexaploid wheat. The extreme structural conservation further validates the strong preservation in the DNA-binding role in the immune signalling mechanism (Aamir et al., 2017). The WRKY domain of the more distant diploid progenitor YrU1 also maintained very high similarity to TaADNR1 (RMSD 0.560 Å) even though this domain is positioned differently in the full-length sequence (residues 1429–1488 for YrU1 vs. 1333-1392 for TaADNR1). This level of conservation supports that the WRKY domain is under high evolutionary conservation and maintenance, suggesting that this is a key functional domain that plays a critical role in immune signalling in wheat species (Sarris et al., 2015). Such a strong conservation of the observed domains is remarkable for NLR proteins which are known for evolving rapidly for pathogen recognition (Jacob et al., 2013; Wang et al., 2020b). However, this high level of conservation may result in the functioning of the WRKY domain being highly susceptible to disruption of its structure and mutations in this domain’s core residues could negatively impact the relay of an immune signal and affect the evolutionary function (Nicolis and Venter, 2018; Wang et al., 2020a).

## Supporting information

Supplementary Figures, Tables and Scripts

## Acknowledgements

This work was funded by public grants from the Winter Cereal Trust (WCT/W/2007/04 Induced systemic resistance in wheat).

## Supplementary information

**Table S1:** The amino acid sequence comparison of the WRKY domain region of TaADNR1 and AtWRKY41. The table lists residue substitutions and their physicochemical classification identities. The substitutions are classified based on the changes in polarity and charge between the residues.

**Table S2:** The amino acid sequence comparison of the WRKY domain region of TaADNR1 and AtWRKY46. The table lists residue substitutions and their physicochemical classification identities. The substitutions are classified based on the changes in polarity and charge between the residues.

**Table S3:** The amino acid sequence comparison of the WRKY domain region of AtWRKY41 and AtWRKY46. The table lists residue substitutions and their physicochemical classification identities. The substitutions are classified based on the changes in polarity and charge between the residues.

**Table S4:** The BLAST^™^ search annotations of the six identified *T. urartu* WRKY transcription factors

**Table S5:** The structural predictions and comparison metrics of the different wheat ANK-NLR-WRKY proteins

**Table S6:** The structural predictions and comparison metrics of the *T. aestivum* TaADNR1 protein WRKY-only domain against the *A. thaliana* AtWRKY41 and AtWRKY46 proteins. The average pLDDT score and high confidence percentages for the three isolated WRKY domains for each protein are indicated, including the RMSD measured in Ångstrom (Å) with reference to the TaADNR1 isolated WRKY domain. The key insight into the overall confidence of the predictions is also indicated.

**Figure S1:** The multiple sequence alignment of the full-length ANK–NLR–WRKY proteins from *T. urartu* (YrU1), *T. dicoccoides* (TRIDC5AG050380), and *T. aestivum* (TaADNR1).

**Figure S2:** The high- and low-confidence percentages of the full-length and WRKY-only proteins from the AlphaFold2 structure predictions of TaADNR1, TRIDC5AG050380 and YrU1

