## Supplementary Figures, Tables and Scripts for "Comparative Analysis of Orthologous ANK-NLR-WRKY Proteins in Wheat Species: Evolutionary Divergence and Structural Characteristics"

**Table S1**: The amino acid sequence comparison of the WRKY domain region of TaADNR1 and AtWRKY41. The table lists residue substitutions and their physicochemical classification identities. The substitutions are classified based on the changes in polarity and charge between the residues.

| Position | Region | TaADNR1 | AtWRKY41 | Exact substitution | TaADNR1 | AtWRKY41 | Physicochemical change |
|---|---|---|---|---|---|---|---|
| **1** | Upstream | E | D | **Glu1Asp** | Polar, negatively charged | Polar, negatively charged | Polar, negatively charged → Polar, negatively charged |
| **3** | Upstream | G | I | **Gly3Ile** | Non-polar | Non-polar | Non-polar → Non-polar |
| **15** | Downstream | I | L | **Ile15Leu** | Non-polar | Non-polar | Non-polar → Non-polar |
| **17** | Downstream | S | A | **Ser17Ala** | Polar, uncharged | Non-polar | **Polar, uncharged → Non-polar** |
| **18** | Downstream | M | K | **Met18Lys** | Non-polar | Polar, positively charged | **Non-polar → Polar, positively charged** |
| **19** | Downstream | H | F | **His19Phe** | Polar, ionisable | Non-polar | **Polar, ionisable → Non-polar** |
| **22** | Downstream | A | S | **Ala22 Ser** | Non-polar | Polar, uncharged | **Non-polar → Polar, uncharged** |
| **24** | Downstream | F | Y | **Phe24Tyr** | Non-polar | Polar, uncharged | **Non-polar → Polar, uncharged** |
| **28** | Zinc-finger | H | F | **His28Phe** | Polar, ionisable | Non-polar | **Polar, ionisable → Non-polar** |
| **30** | Zinc-finger | H | N | **His30Asn** | Polar, ionisable | Polar, uncharged | Polar, ionisable → Polar, uncharged |
| **31** | Zinc-finger | V | T | **Val31Thr** | Non-polar | Polar, uncharged | **Non-polar → Polar, uncharged** |
| **32** | Zinc-finger | K | Q | **Lys32Gln** | Polar, positively charged | Polar, uncharged | Polar, positively charged → Polar, uncharged |
| **33** | Zinc-finger | G | Y | **Gly33Tyr** | Non-polar | Polar, uncharged | **Non-polar → Polar, uncharged** |
| **35** | Downstream | P | W | **Pro35Trp** | Non-polar | Non-polar | Non-polar → Non-polar |
| **36** | Downstream | V | A | **Val36Ala** | Non-polar | Non-polar | Non-polar → Non-polar |
| **43** | Downstream | T | S | **Thr43Ser** | Polar, uncharged | Polar, uncharged | Polar, uncharged → Polar, uncharged |
| **44** | Downstream | S | D | **Ser44Asp** | Polar, uncharged | Polar, negatively charged | Polar, uncharged → Polar, negatively charged |
| **45** | Downstream | T | G | **Thr45Gly** | Polar, uncharged | Non-polar | **Polar, uncharged → Non-polar** |
| **48** | Downstream | L | T | **Leu48Thr** | Non-polar | Polar, uncharged | **Non-polar → Polar, uncharged** |
| **49** | Downstream | L | I | **Leu49Ile** | Non-polar | Non-polar | Non-polar → Non-polar |
| **51** | Downstream | D | E | **Asp51Glu** | Polar, negatively charged | Polar, negatively charged | Polar, negatively charged → Polar, negatively charged |
| **53** | Downstream | V | T | **Val53Thr** | Non-polar | Polar, uncharged | **Non-polar → Polar, uncharged** |
| **55** | Downstream | H | R | **His55Arg** | Polar, ionisable | Polar, positively charged | Polar, ionisable → Polar, positively charged |
| **57** | Downstream | E | T | **Glu57Thr** | Polar, negatively charged | Polar, uncharged | Polar, negatively charged → Polar, uncharged |

**Table S2**: The amino acid sequence comparison of the WRKY domain region of TaADNR1 and AtWRKY46. The table lists residue substitutions and their physicochemical classification identities. The substitutions are classified based on the changes in polarity and charge between the residues.

| **Position** | **Region** | **TaADNR1** | **AtWRKY46** | **Exact substitution** | **TaADNR1** | **AtWRKY46** | **Physicochemical change** |
|---|---|---|---|---|---|---|---|
| **1** | Upstream | E | D | **Glu1Asp** | Polar, negatively charged | Polar, negatively charged | Polar, negatively charged → Polar, negatively charged |
| **4** | Upstream | F | H | **Phe4 His** | Non-polar | Polar, ionisable | **Non-polar → Polar, ionisable** |
| **5** | Upstream | S | C | **Ser5Cys** | Polar, uncharged | Polar, uncharged | Polar uncharged → Polar, uncharged |
| **13** | Downstream | D | E | **Asp13Glu** | Polar, negatively charged | Polar, negatively charged | Polar, negatively charged → Polar, negatively charged |
| **15** | Downstream | I | H | **Ile15His** | Non-polar | Polar, ionisable | **Non-polar → Polar, ionisable** |
| **18** | Downstream | M | K | **Met18Lys** | Non-polar | Polar, positively charged | **Non-polar → Polar, positively charged** |
| **19** | Downstream | H | N | **His19Asn** | Polar, ionisable | Polar, uncharged | Polar, ionisable → Polar, uncharged |
| **24** | Downstream | F | Y | **Phe24Trr** | Non-polar | Polar, uncharged | **Non-polar → Polar, uncharged** |
| **30** | Zinc-finger | H | F | **His30Phe** | Polar, ionisable | Non-polar | **Polar, ionisable → Non-polar** |
| **31** | Zinc-finger | V | T | **Val31Thr** | Non-polar | Polar, uncharged | **Non-polar → Polar, uncharged** |
| **32** | Zinc-finger | K | Q | **Lys32Gln** | Polar, positively charged | Polar, uncharged | Polar, positively charged → Polar, uncharged |
| **33** | Zinc-finger | G | D | **Gly33Asp** | Non-polar | Polar, negatively charged | **Non-polar → Polar, negatively charged** |
| **35** | Downstream | P | L | **P 35 Leu** | Non-polar | Non-polar | Non-polar → Non-polar |
| **36** | Downstream | V | A | **Val36Ala** | Non-polar | Non-polar | Non-polar → Non-polar |
| **37** | Downstream | T | V | **Thr37Val** | Polar, uncharged | Non-polar | **Polar, uncharged → Non-polar** |
| **42** | Downstream | R | K | **Arg42Lys** | Polar, positively charged | Polar, positively charged | Polar, positively charged → Polar, positively charged |
| **43** | Downstream | T | S | **Thr43Ser** | Polar, uncharged | Polar, uncharged | Polar, uncharged → Polar, uncharged |
| **44** | Downstream | S | D | **Ser44Asp** | Polar, uncharged | Polar, negatively charged | Polar, uncharged → Polar, negatively charged |
| **48** | Downstream | L | S | **Leu48Ser** | Non-polar | Polar, uncharged | **Non-polar → Polar, uncharged** |
| **51** | Downstream | D | E | **Asp51Glu** | Polar, negatively charged | Polar, negatively charged | Polar, negatively charged → Polar, negatively charged |
| **53** | Downstream | V | K | **Val53Lys** | Non-polar | Polar, positively charged | **Non-polar → Polar, positively charged** |
| **55** | Downstream | H | L | **His55 Leu** | Polar, ionisable | Non-polar | **Polar, ionisable → Non-polar** |
| **57** | Downstream | E | N | **Glu57Asn** | Polar, negatively charged | Polar, uncharged | Polar, negatively charged → Polar, uncharged |

**Table S3**: The amino acid sequence comparison of the WRKY domain region of AtWRKY41 and AtWRKY46. The table lists residue substitutions and their physicochemical classification identities. The substitutions are classified based on the changes in polarity and charge between the residues.

| Position | Region | AtWRKY41 | AtWRKY46 | Exact substitution | AtWRKY41 | AtWRKY46 | Physicochemical change |
|---|---|---|---|---|---|---|---|
| **3** | Upstream | I | G | **Ile3Gly** | Non-polar | Non-polar | Non-polar → Non-polar |
| **4** | Upstream | F | H | **Phe4His** | Non-polar | Polar, ionisable | **Non-polar → Polar, ionisable** |
| **5** | Upstream | S | C | **Ser5Cys** | Polar, uncharged | Polar, uncharged | Polar, uncharged → Polar, uncharged |
| **13** | Downstream | D | E | **Asp13Glu** | Polar, negatively charged | Polar, negatively charged | Polar, negatively charged → Polar, negatively charged |
| **15** | Downstream | L | H | **Leu15His** | Non-polar | Polar, ionisable | **Non-polar → Polar, ionisable** |
| **17** | Downstream | A | S | **Ala17Ser** | Non-polar | Polar, uncharged | **Non-polar → Polar, uncharged** |
| **19** | Downstream | F | N | **Phe19Asn** | Non-polar | Polar, uncharged | **Non-polar → Polar, uncharged** |
| **22** | Downstream | S | A | **Ser22Ala** | Polar, uncharged | Non-polar | **Polar, uncharged → Non-polar** |
| **28** | Zinc-finger | F | H | **Phe28His** | Non-polar | Polar, ionisable | **Non-polar → Polar, ionisable** |
| **30** | Zinc-finger | N | F | **Asn30Phe** | Polar, uncharged | Non-polar | **Polar, uncharged → Non-polar** |
| **33** | Zinc-finger | Y | D | **Tyr33Asp** | Polar, uncharged | Polar, negatively charged | Polar, uncharged → Polar, negatively charged |
| **35** | Downstream | W | L | **Trp35 Leu** | Non-polar | Non-polar | Non-polar → Non-polar |
| **37** | Downstream | T | V | **Thr37Val** | Polar, uncharged | Non-polar | **Polar, uncharged → Non-Polar** |
| **42** | Downstream | R | K | **Arg42Lys** | Polar, positively charged | Polar, positively charged | Polar, positively charged → Polar, positively charged |
| **45** | Downstream | G | T | **Gly45Thr** | Non-polar | Polar, uncharged | **Non-polar → Polar, uncharged** |
| **48** | Downstream | T | S | **Thr48Ser** | Polar, uncharged | Polar, uncharged | Polar, uncharged → Polar, uncharged |
| **49** | Downstream | I | L | **Ile49 Leu** | Non-polar | Non-polar | Non-polar → Non-polar |
| **53** | Downstream | T | K | **Thr53Lys** | Polar, uncharged | Polar, positively charged | Polar, uncharged → Polar, positively charged |
| **55** | Downstream | R | L | **Arg55 Leu** | Polar, positively charged | Non-polar | **Polar, positively charged → Non-polar** |
| **57** | Downstream | T | N | **Thr57Asn** | Polar, uncharged | Polar, uncharged | Polar, uncharged → Polar, uncharged |

**Table S4**: The BLAST™ search annotations of the six identified *T. urartu* WRKY transcription factors

| *Triticum urartu* WRKY TFs | Top Arabidopsis BLAST hit (TFDB) | Score (TFDB) | E-value (TFDB) | Top Arabidopsis BLAST hit (TAIR) | Score (Bits) (TAIR) | E-value (TAIR) | Interpretation | Function |
|---|---|---|---|---|---|---|---|---|
| **TuG1812G0100004483.01** | AtWRKY41 | 270 | 3e-29 | AtWRKY41 | 108 | 7e-28 | WRKY41-like | Defense signal, fungal resistance transduction, seed dormancy and anthocyanin biosynthesis |
| **TuG1812G0300002773.01** | AtWRKY41 | 338 | 6e-39 | AtWRKY41 | 134 | 2e-37 | WRKY41-like | Defense signal, fungal resistance transduction, seed dormancy and anthocyanin biosynthesis |
| **TuG1812S0003165700.01** | AtWRKY41 | 302 | 8e-35 | AtWRKY41 | 120 | 2e-33 | WRKY41-like | Defense signal, fungal resistance transduction, seed dormancy and anthocyanin biosynthesis |
| **TuG1812G0500002157.01** | AtWRKY46 | 303 | 4e-33 | AtWRKY46 | 121 | 1e-31 | WRKY46-like | Salicylic acid synthesis upon pathogen attack, drought and salt stress, root-to-shoot iron translocation |
| **TuG1812G0200001696.01** | AtWRKY41 | 296 | 1e-32 | AtWRKY41 | 118 | 2e-31 | WRKY41-like | Defense signal, fungal resistance transduction, seed dormancy and anthocyanin biosynthesis |

**Table S5**: The structural predictions and comparison metrics of the different wheat ANK-NLR-WRKY proteins

| Protein | Source | Full-length Avg. pLDDT | WRKY domain Avg. pLDDT | Full-length High-Confidence Residues (70-90%) | WRKY domain High-Confidence Residues (70-90%) | Vs. TaADNR1 RMSD (Full-Length, Å) | Vs. TaADNR1 RMSD (WRKY domain, Å) | Key Structural Insight |
|---|---|---|---|---|---|---|---|---|
| **TaADNR1** | *T. aestivum* | 71.91 | 82.27 | 53.6 | 80 | Reference | Reference | High confidence (suitable for comparative structural analysis) |
| **YrU1** | *T. urartu* | 68.32 | 78.48 | 52.1 | 76.7 | 0.746 | 0.56 | Medium confidence (suitable for fold-level analysis) |
| **TRIDC5AG050380** | *T. dicoccoides* | 71.64 | 81.61 | 53.9 | 81.7 | 0.424 | 0.13 | High confidence (suitable for comparative structural analysis) |

*Note: pLDDT - per-residue Predicted Local Distance Difference Test. RMSD - root mean square deviation.

**Table S6**: The structural predictions and comparison metrics of the *T. aestivum* TaADNR1 protein WRKY-only domain against the *A. thaliana* AtWRKY41 and AtWRKY46 proteins. The average pLDDT score and high confidence percentages for the three isolated WRKY domains for each protein are indicated, including the RMSD measured in Ångstrom (Å) with reference to the TaADNR1 isolated WRKY domain. The key insight into the overall confidence of the predictions is also indicated.

| Protein | Source | WRKY domain Avg. pLDDT | WRKY domain High-Confidence Residues (%) | Vs. TaADNR1 RMSD (WRKY domain) | Key Structural Insight |
|---|---|---|---|---|---|
| **TaADNR1** | *T. aestivum* | 82.27 | 80 (70-90%) (48 residues) | Reference | High confidence (suitable for comparative structural analysis) |
| **AtWRKY41** | *A. thaliana* | 93.71 | 88.3 (>90%) (53 residues) | 0.360 | Very high confidence (suitable for comparative structural analysis) |
| **AtWRKY46** | *A. thaliana* | 92.34 | 78.7 (>90%) (48 residues) | 0.378 | Very high confidence (suitable for comparative structural analysis) |

*Note: Å - measurement in Ångstrom. pLDDT - per-residue Predicted Local Distance Difference Test. RMSD - root mean square deviation.

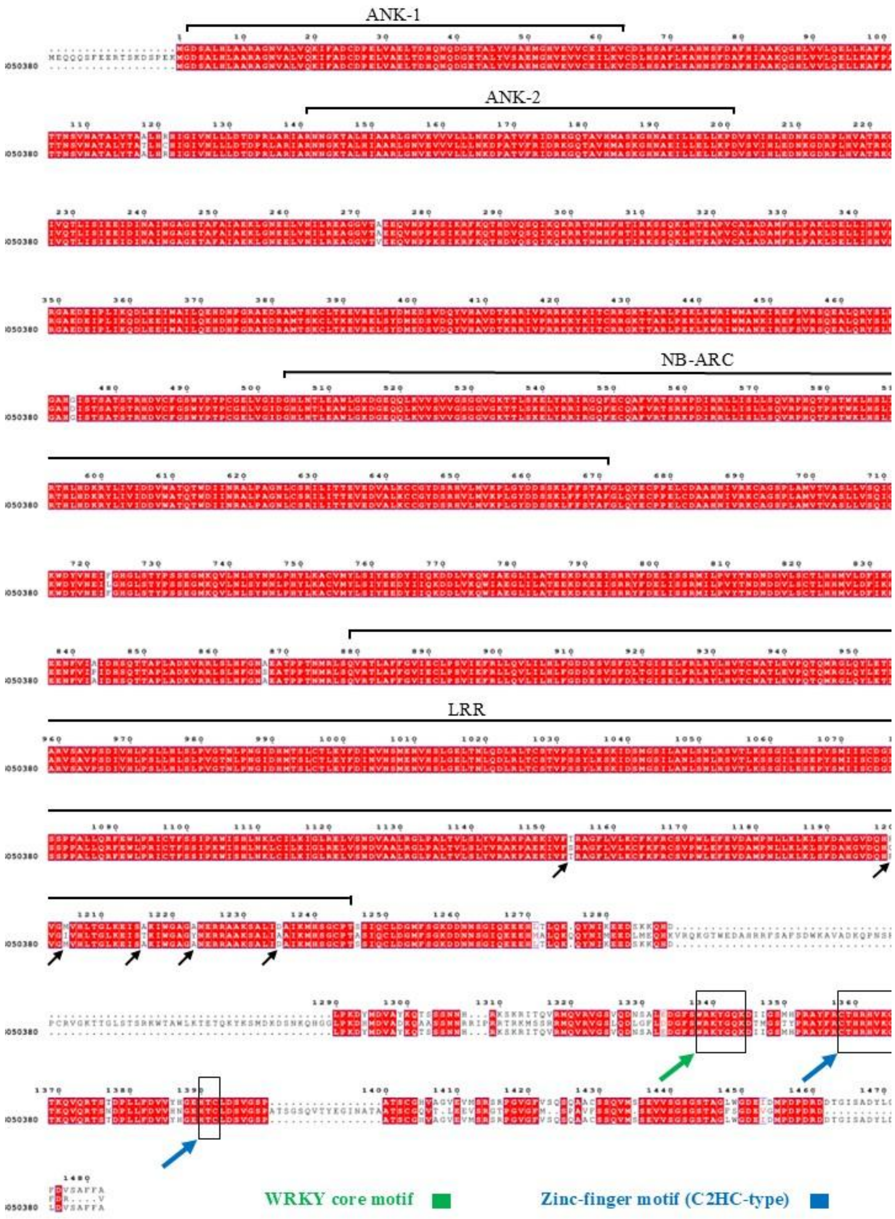


**Figure S1**: The multiple sequence alignment of the full-length ANK–NLR–WRKY proteins from *T. urartu* (YrU1), *T. dicoccoides* (TRIDC5AG050380), and *T. aestivum* (TaADNR1).

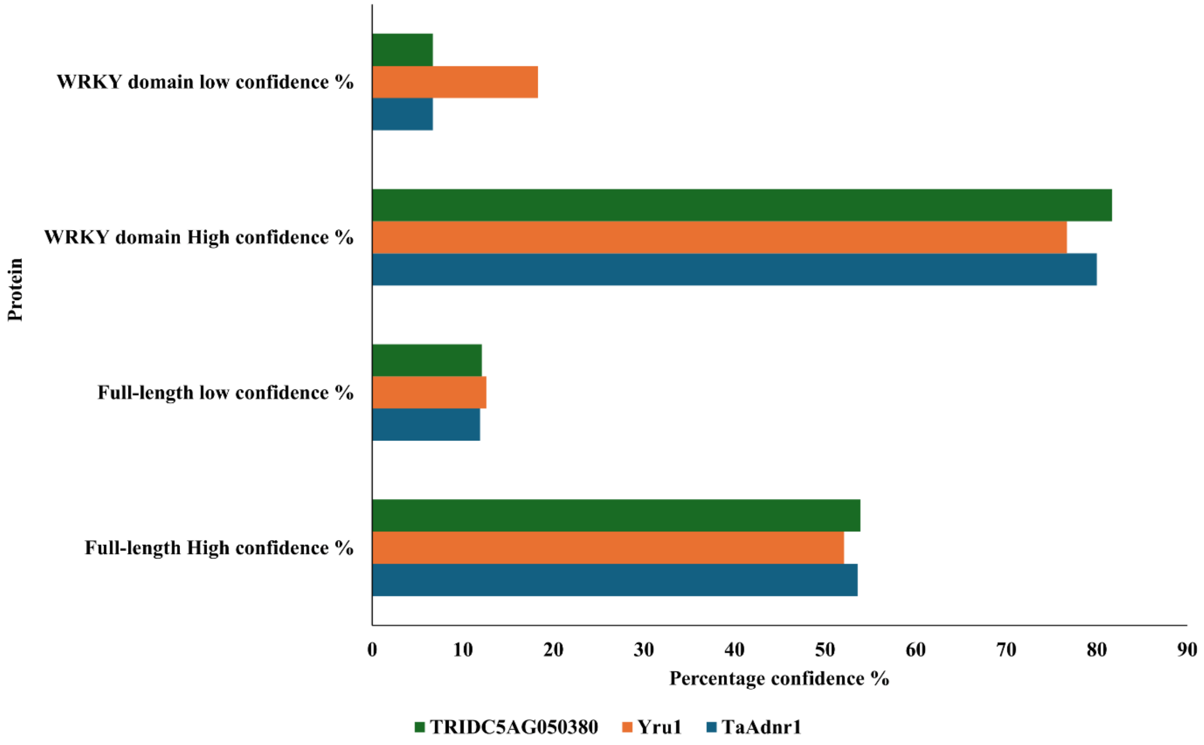


**Figure S2**: The high- and low-confidence percentages of the full-length and WRKY-only proteins from the AlphaFold2 structure predictions of TaADNR1, TRIDC5AG050380 and YrU1

**The script for WRKY Domain pLDDT extraction (get_plddt_WRKY.py)**
This following script was used to extract the per-residue pLDDT scores from the AlphaFold2 .pkl output files, to analyse the WRKY domain statistical data, and to compare the full-length protein data:

```
"""
get_plddt_WRKY.py - WRKY Domain pLDDT Calculator for AlphaFold2 Outputs

This script extracts pLDDT scores from AlphaFold2 .pkl result files,
calculates statistics for user-specified WRKY domains, and compares
domain confidence with full-length protein metrics.

Usage: python get_plddt_WRKY.py
"""

import pickle
import numpy as np
import os

print("="*60)
print("AlphaFold WRKY Domain pLDDT Calculator")
print("="*60)

# ===========================================================================
# STEP 1: Get the .pkl file path from the user
# ===========================================================================
print("\nSTEP 1: Locate your .pkl file.")
print("Example: C:\\Users\\biotechlab\\Documents\\YrU1\\result_model_1_pred_0.pkl")
file_path = input("Paste the FULL path to your .pkl file: ").strip('\"\' ')

# Check if file exists
if not os.path.exists(file_path):
    print(f"\n❌ ERROR: File not found at: {file_path}")
    input("Press Enter to exit...")
    exit()

# ===========================================================================
# STEP 2: Get the WRKY domain residue range from the user
# ===========================================================================
print("\nSTEP 2: Define the WRKY domain residue range.")
print("Example for TaAdnr1: 1333 1392")
print("Example for YrU1: 1429 1488")

try:
    start_res = int(input("Enter START residue number: "))
    end_res = int(input("Enter END residue number: "))
except ValueError:
    print("❌ Please enter valid numbers.")
    input("Press Enter to exit...")
    exit()

if start_res >= end_res:
    print("❌ Start residue must be less than end residue.")
    input("Press Enter to exit...")
    exit()
```

```
# ====================================================================
# STEP 3: Load data and extract pLDDT for the domain
# ====================================================================
print("\n" + "="*60)
print("ANALYZING...")
print("="*60)

try:
    # Load the .pkl file
    with open(file_path, 'rb') as f:
        data = pickle.load(f)

    # Find the pLDDT array (trying common keys)
    plddt_full = None
    for key in ['plddt', 'confidence', 'confidence_score']:
        if key in data:
            plddt_full = np.array(data[key], dtype=np.float32)
            print(f"✓ Found pLDDT array using key: '{key}'")
            print(f"✓ Full protein length: {len(plddt_full)} residues")
            break

    if plddt_full is None:
        print("❌ Could not find pLDDT data in the file.")
        print("Available keys:", list(data.keys()))
        input("Press Enter to exit...")
        exit()

    # Extract WRKY domain region (0-indexed array, 1-indexed residues)
    start_idx = start_res - 1
    end_idx = end_res  # Python slicing excludes the last index

    if start_idx < 0 or end_idx > len(plddt_full):
        print(f"❌ Error: Domain range {start_res}-{end_res} is outside protein range (1-{len(plddt_full)})")
        input("Press Enter to exit...")
        exit()

    plddt_domain = plddt_full[start_idx:end_idx]
    domain_length = end_res - start_res + 1

    print(f"\n✓ WRKY domain extracted: residues {start_res}-{end_res}")
    print(f"✓ Domain length: {domain_length} residues")

    # ====================================================================
    # STEP 4: Calculate statistics and display results
    # ====================================================================
    print("\n" + "="*60)
    print("RESULTS FOR WRKY DOMAIN")
    print("="*60)

    print(f"Protein file: {os.path.basename(file_path)}")
    print(f"WRKY domain: residues {start_res}-{end_res}")
    print(f"Domain length: {domain_length} residues")

    print(f"\n📊 CONFIDENCE SCORES:")
    print(f"  • Average pLDDT: {np.mean(plddt_domain):.2f}")
```

```
    print(f"  • Maximum pLDDT: {np.max(plddt_domain):.2f}")
    print(f"  • Minimum pLDDT: {np.min(plddt_domain):.2f}")
    print(f"  • Standard deviation: {np.std(plddt_domain):.2f}")

    # Compare with whole protein
    whole_avg = np.mean(plddt_full)
    domain_avg = np.mean(plddt_domain)
    diff = domain_avg - whole_avg

    print(f"\n🔍 COMPARISON WITH WHOLE PROTEIN:")
    print(f"  • Whole protein average pLDDT: {whole_avg:.2f}")
    print(f"  • WRKY domain average pLDDT: {domain_avg:.2f}")
    print(f"  • Difference (Domain - Whole): {diff:+.2f}")

    # Confidence band analysis for the domain
    print(f"\n🎯 CONFIDENCE DISTRIBUTION IN WRKY DOMAIN:")

    bands = [
        ('Very high (>90)', plddt_domain > 90),
        ('High (70-90)', (plddt_domain >= 70) & (plddt_domain <= 90)),
        ('Low (50-70)', (plddt_domain >= 50) & (plddt_domain < 70)),
        ('Very low (<50)', plddt_domain < 50)
    ]

    for name, condition in bands:
        count = np.sum(condition)
        percentage = (count / len(plddt_domain)) * 100
        print(f"  • {name}: {count:3d} residues ({percentage:5.1f}%)")

    # Interpretation
    print(f"\n💡 INTERPRETATION:")
    if domain_avg > 90:
        print(f"  • WRKY domain: VERY HIGH confidence (suitable for detailed mechanistic analysis)")
    elif domain_avg > 80:
        print(f"  • WRKY domain: HIGH confidence (suitable for comparative structural analysis)")
    elif domain_avg > 70:
        print(f"  • WRKY domain: MEDIUM confidence (suitable for fold-level analysis)")
    elif domain_avg > 50:
        print(f"  • WRKY domain: LOW confidence (interpret with caution)")
    else:
        print(f"  • WRKY domain: VERY LOW confidence (unreliable for structural analysis)")

    if diff > 5:
        print(f"  • The WRKY domain is SIGNIFICANTLY MORE CONFIDENT than the whole protein")
    elif diff > 2:
        print(f"  • The WRKY domain is MORE CONFIDENT than the whole protein")
    elif diff < -5:
        print(f"  • Warning: The WRKY domain is LESS CONFIDENT than the whole protein")

    print("\n" + "="*60)

except Exception as e:
    print(f"\n❌ An error occurred: {str(e)}")
    import traceback
    traceback.print_exc()
```

```
input("\nPress Enter to exit...")
```
